# Combined *in situ* hybridization chain reaction and immunostaining to visualize gene expression in whole-mount *Drosophila* central nervous systems

**DOI:** 10.1101/2021.08.02.454831

**Authors:** Julia C. Duckhorn, Ian Junker, Yun Ding, Troy R. Shirangi

**Affiliations:** Department of Biology, Villanova University, Villanova, PA 19312; Department of Biology, University of Pennsylvania, Philadelphia, PA 19104

## Abstract

Methods to visualize gene expression in the *Drosophila* central nervous system are important in fly neurogenetic studies. In this chapter, we describe a detailed protocol that sequentially combines *in situ* hybridization chain reaction (HCR) and immunostaining to detect mRNA and protein expression in whole-mount *Drosophila* larval and adult central nervous systems. We demonstrate the application of *in situ* HCR in comparisons of nervous system gene expression between *Drosophila* species, and in the validation of single-cell RNA-Seq results in the fly nervous system. Our protocol provides a simple, robust, multiplexable, and relatively affordable means to quantitatively visualize gene expression in the nervous system of flies, facilitating its general use in fly neurogenetic studies.

## 1. INTRODUCTION

Neurogenetic studies of *Drosophila* often require methods to visualize and quantify gene transcription in the larval or adult central nervous system. Fluorescence *in situ* hybridization of mRNAs, or mRNA-FISH, is widely used across animal systems to detect gene expression in a variety of whole-mount tissue-types. However, the application of mRNA-FISH on whole-mount *Drosophila* central nervous systems has faced long-standing challenges, including weak probe penetration and poor sample preservation, especially in adult nervous systems.

*In situ* hybridization chain reaction (HCR) is an enzyme-free method to detect one or more sequence-specific nucleic acids in cell, tissue, and whole-mount samples via detection and amplification stages [1–4]. During detection, an initiator sequence assembles on target mRNA molecules using adjacent, paired DNA probes. The initiator sequence subsequently triggers the self-assembly of a fluorescently tagged DNA polymer from a mixture of two amplifier hairpins. The use of short DNA probe-sets greatly enhances tissue penetration and signal detection. The detection and amplification reactions are done under mild experimental conditions, thus improving sample integrity and permitting the application of immunostaining protocols. Moreover, the use of split-paired probes suppresses off-target amplification, thereby facilitating high signal-to-noise ratios [4].

In recent years, two whole-mount mRNA-FISH protocols have been developed for use on *Drosophila* central nervous systems [5, 6]. Both protocols detect mRNA expression with single-molecule resolution using fluorescently labeled probe-sets but differ in sample preparation and imaging protocols. In comparison to these protocols, an HCR-based approach is, in principle, more sensitive and robust, allowing it to detect transcripts across different expression levels. It is also more cost-effective and does not require the experimenter to design or create probe-sets, which are provided by Molecular Instruments™. HCR also offers greater flexibility to mix and match color channels for multiplexing and post-HCR immunostaining, since the amplifier hairpins and not the probes themselves are labeled with a fluorochrome. Although *in situ* HCR-based FISH has been successfully implemented using whole-mount *Drosophila* CNS tissues [7, 8], the procedure is not widely used in the field.

In this chapter, we report a detailed *in situ* HCR protocol that provides a multiplexable, straightforward, affordable, rapid, sensitive, and robust means to detect mRNA expression in the *Drosophila* larval and adult central nervous system using confocal light microscopy. Our protocol can be used with standard immunohistochemistry protocols to simultaneously detect protein expression. We demonstrate the utility of *in situ* HCR with two important applications. First, we visualize the mRNA expression of multiple orthologous genes in the adult and larval central nervous systems of two *Drosophila* species. Second, we validate recently published single-cell RNA-Seq data [9] on neurons derived from a common neuronal stem cell in the fly ventral nervous cord.

## 2. MATERIALS

### 2.1 Split-Initiator Probes and Amplification Hairpins

Single-stranded DNA HCR probe-sets and fluorescently tagged DNA HCR amplifier hairpins were purchased from Molecular Instruments™. Each probe- and hairpin-set are matched by the identity of the HCR initiator. For example, if a particular probe-set utilizes a B1 HCR initiator, a B1 complementary hairpin set is required for successful polymerization. We used a probe-set size of 20 for all *in situ* HCR experiments. Probes are complementary DNA sequences of the desired endogenous mRNA target. The sequences of each probe within a probe-set are available upon request from Molecular Instruments™. The lot numbers for the probe-sets used in this study are: PRH303 (*Antp*), PRE237 (*Ubx*), PRF810 (*Abd-B*), and PR1675 (*Sox21B*). For long term storage, probes and hairpins were aliquoted and frozen at −20°C. During use, aliquots were defrosted and stored short term at 4°C. All fluorescently tagged hairpins were protected from light during storage.

### 2.2 Reagents and Buffers for HCR and Immunostaining

Central nervous systems were dissected in 1X phosphate buffered saline (PBS; Fisher Cat. # BP2438-4) and fixed in 4% paraformaldehyde (PFA; Electron Microscopy Sciences, Cat. # 15710) buffered in 1X PBS. 4% PFA was stored at 4°C for no more than two weeks or as single-use aliquot at −20°C. Probe hybridization buffer (stored at −20°C), probe wash buffer (stored at −20°C), and amplification buffer (stored at 4°C) were purchased from Molecular Instruments™, but can be easily prepared in the lab [4]. 5X SSCT (UltraPure 20X SSC, ThermoFisher Scientific, Cat. # 15557044 plus Tween 20 diluted to 0.1%) was made fresh the day of usage. Nervous tissue was washed in PBTx (1X PBS with 1% Triton X-100) stored at room temperature or 4°C. Nervous systems were blocked in 5% heat-inactivated normal goat serum (NGS; Jackson Immunochemicals, Cat. # 005-000-121) in PBTx. The following primary antibodies were used in this study: mouse anti-Antp 8C11 (Developmental Studies Hybridoma Bank, diluted 1:20 in PBTx/5% NGS), mouse anti-Ubx FP3.38 (DSHB, diluted 1:20), and mouse anti-Abd-B 1A2E9 (DSHB, diluted 1:10). All primary antibodies were used with a goat anti-mouse Alexa Fluor® 568 secondary diluted in PBTx/5% NGS (Invitrogen, Cat. # A11004, diluted 1:500).

### 2.3 Reagents for mounting

We mounted larval and adult nervous systems in VECTASHIELD (Cat. # H-1000) or ProLong Gold antifade with DAPI (Invitrogen, Cat. # P36931). If users prefer to clear with xylenes before mounting, we recommend passing the nervous systems through an ascending ethanol series (30%, 50%, 70%, 95%, 100%, 100%) for 5 minutes each, then placing the coverslip-adhered (Corning; Cat. # 2845-22) nervous systems in xylenes (Sigma-Aldrich, Cat. # 247642-500ML) twice for 5 minutes. The coverslip with nervous systems adhered to it can be mounted on a slide using Dibutylphthalate Polystyrene Xylene (DPX; Sigma-Aldrich, Cat. # 06522-500ML).

### 2.4 Drosophila strains

Fly stocks were maintained at room temperature. The following genotypes were used: *D. melanogaster* Canton S and *D. yakuba* Ivory Coast (UCSD Stock #14021-0261.02).

### 2.5 Confocal imaging

Central nervous systems were imaged on Leica SP8 (larval tissues) and Leica DMi8 (adult tissues) confocal microscopes at 40X and 63X with optical sections at 0.3 or 1 μm intervals.

## 3. METHODS

### 3.1 Multiplexable HCR

To demonstrate the use of *in situ* HCR on the whole-mount central nervous system of *Drosophila melanogaster* (Canton S), we simultaneously probed the mRNA expression of three *Hox* genes in larvae and adults using the protocol described below: *Antennapedia* (*Antp*), *Ultrabithorax* (*Ubx*), and *Abdominal-B* (*Abd-B*). The expression of these *Hox* genes has been well-characterized in the fly nervous system [9], and good antibodies are available against proteins of all three genes. Probe-sets and corresponding amplification hairpins were designed and supplied by Molecular Instruments™ (Antp-B1/B1-Alexa Fluor® 488, Ubx-B2/B2-Alexa Fluor® 594, and Abd-B-B3/B3-Alexa Fluor® 647). The HCR protocol takes three days to complete. An overview of the protocol workflow is illustrated in **Fig. 1**. Exemplary images of resulting larval and adult nervous systems are shown in **Fig. 2**.

**Figure 1.**
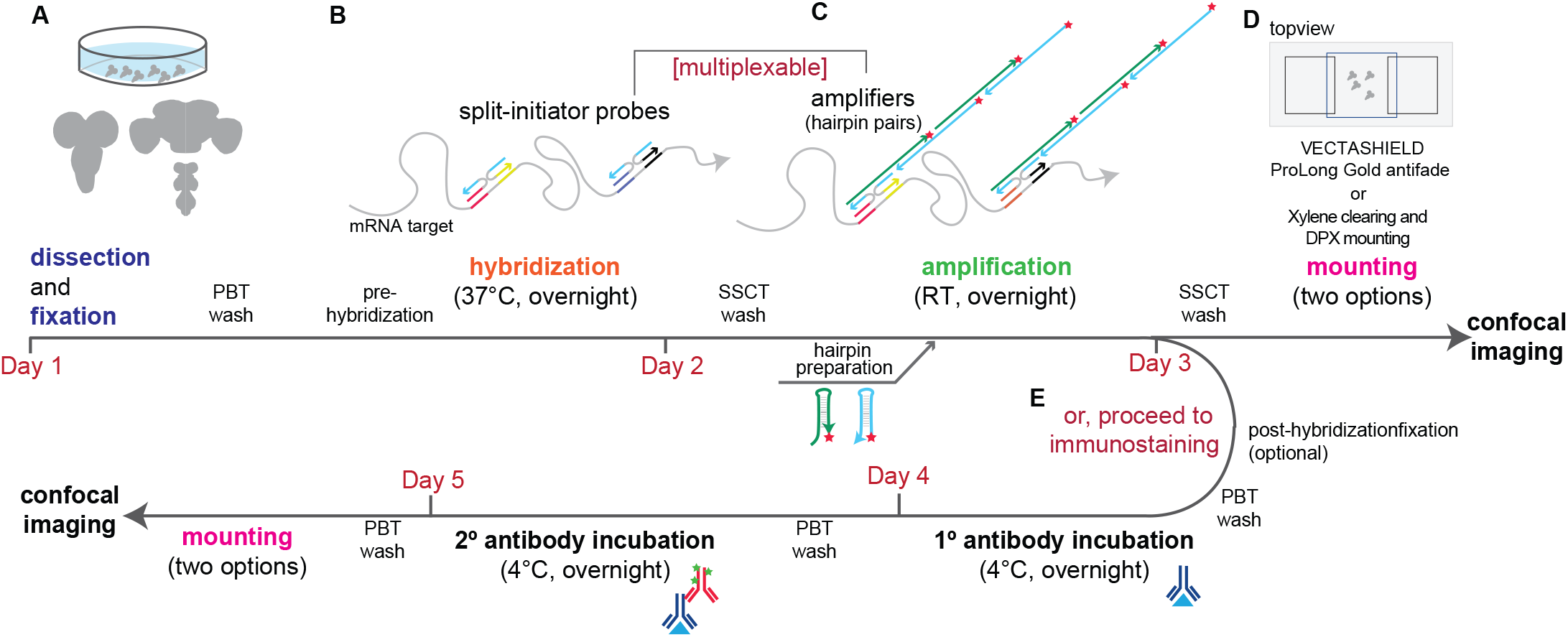
Overview of the *in situ* HCR protocol combined with immunostaining. Summary of the main steps of *in situ* HCR and the mechanisms of detection and amplification are provided along a timeline. (A) Dissection and fixation. All steps are performed in a glass dish. (B) HCR hybridization. mRNA targets are bounded by split-initiator probe-sets (we used a set of 20 pairs in this study). (C) HCR amplification. mRNA targets are illuminated in the amplification stage, where initiators assembled by individual probe pairs trigger the self-assembly of hairpin amplifiers conjugated with fluorophores (denoted by stars). HCR is multiplexable by matching probe-sets designed for different transcripts with hairpin pairs conjugated with different fluorophores. The timeline is the same regardless of the number of targeted mRNA. (D) Mounting. Samples can be mounted directly with generic mounting media such as VECTASHIELD or ProLong Gold Antifade or be subjected to xylene clearing and DPX mounting for potentially optimized signal-to-noise ratio. (E) Immunostaining. Users may perform this step after HCR and an optional post-HCR PFA fixation (for better preservation of HCR signals). Some antibodies are sensitive to this additional fixation step, and users can skip it or adjust the conditions for a more desirable outcome. Users may also try immunostaining before HCR, if the antibody staining is consistently problematic.

**Figure 2.**
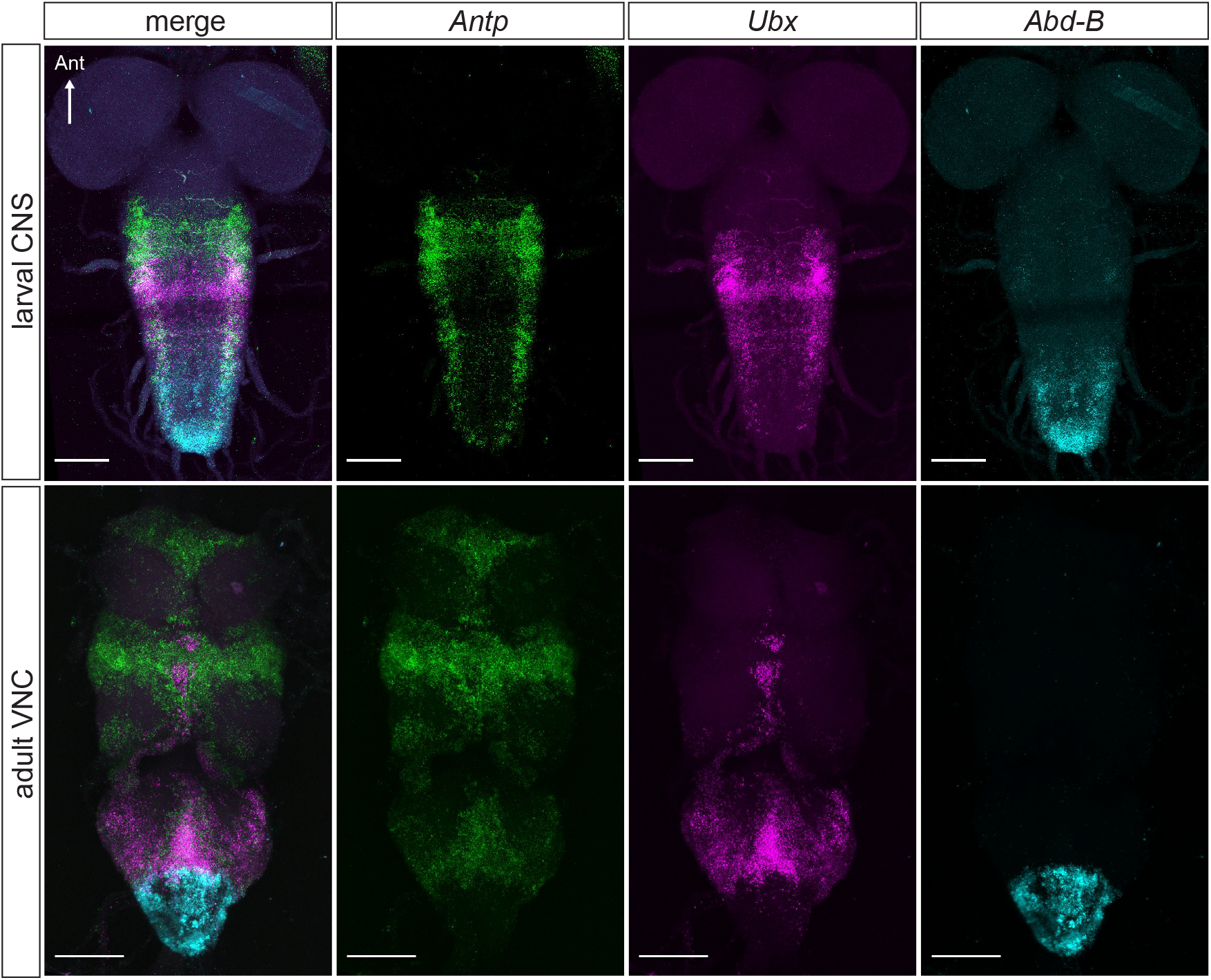
Detection of *Hox* gene mRNA expression in the larval and adult central nervous system of *D. melanogaster* by *in situ* HCR. Antp-B1, Ubx-B2, and Abd-B-B3 probe-sets were used with B1-Alexa Fluor® 488, B2-Alexa Fluor® 594, and B3-Alexa Fluor® 647 amplifiers in multiplexed *in situ* HCR. mRNAs of *Antp* and *Ubx* are expressed primarily in cells of the thoracic neuromeres; *Abd-B* is expressed in cells of the abdominal ganglion. *Hox* gene expression occurs in a pattern consistent with that previously published (Allen *et al*., 2020). Scale bar = 50 μm.

#### Day 1

1. Dissect nervous systems from late-stage 3^rd^ instar larvae or adult female flies aged 4–7 days in cold 1X PBS and transfer them to a glass embryo dish (Electron Microscopy Sciences; Cat. # 70543-30) on ice containing 1X PBS. All subsequent steps are performed in the dish.
2. Fix the nervous systems in 4% PFA fixation buffer for 35 minutes at room temperature. Following fixation, quickly rinse samples three times with PBTx, then wash the samples with PBTx four times for 5 minutes each at room temperature.
3. Pre-hybridize samples in 200 μl prewarmed probe hybridization buffer for 30 minutes at 37°C. In case the tissues are floating on the surface of the buffer, gently prod the tissues downward so they become immersed in the buffer. During pre-hybridization, prepare the probe solution by adding 0.8 pmol of each probe-set to 200 μl of pre-warmed probe hybridization buffer.
4. Replace the pre-hybridization solution with the probe solution, and incubate overnight (12–16 hours) at 37°C. The embryo dish should be covered and sealed tightly with parafilm to prevent the probe solution from evaporating overnight. All following steps should be completed with protection from light.

#### Day 2

5. Pre-heat the probe wash buffer for 15 minutes at 37°C. Remove probe solution and wash the nervous systems with 200 μl of pre-heated probe wash buffer four times, 15 minutes each at 37°C. Next, remove the probe wash buffer and wash the tissues with 5X SSCT three times, 5 minutes at room temperature. 5X SSCT should be prepared fresh the day of use.
6. Pre-amplify the samples in 200 μl of amplification buffer for 30 minutes at room temperature. Prewarm the amplification buffer to room temperature before use.
7. During pre-amplification, prepare the hairpin solution: Each sample receives 12 pmol of each hairpin in a total of 200 μl of amplification buffer. Add 12 pmol of hairpin h1 to one Eppendorf tube and 12 pmol of hairpin h2 to another Eppendorf tube (*i.e*., if the stock solution of each hairpin is 3 μM or 3 pmol/μl, the experimenter should add 4 μl of the hairpin to the tube). Snap-cool by placing both tubes in a preheated heat-block set at 95°C for 90 seconds, then allow the hairpins to cool to room temperature for 30 minutes in a dark drawer. Then add 100 μl of amplification buffer to each hairpin, mix gently, and combine the two hairpins for a total of 200 μl.
8. Replace pre-amplification solution with the hairpin solution and incubate overnight (12–16 hours) at room temperature. The dish should be covered and sealed with parafilm.

#### Day 3

9. Remove the hairpin solution and wash samples with 5X SSCT at room temperature using the following series: two times, 5 minutes each; two times, 30 minutes each; and one time for 5 minutes.
10. Mount the nervous systems. Users may mount tissues using standard mounting procedures with VECTASHIELD or ProLong Gold antifade. Alternatively, users may choose to mount with DPX after clearing the nervous systems with xylenes. Users should note that compared to tissues that pass-through immunohistochemistry-only protocols, post-HCR tissues often do not stick well to poly-L-lysine-coated coverslips. This can lead to sample loss in the subsequent ethanol dehydration steps. We therefore recommend that ethanol dehydration be performed in an Eppendorf tube instead of on a poly-L-lysine-coated coverslip. Upon completion of the series, transfer the nervous systems onto a poly-L-lysine-coated coverslip before clearing with xylene, which makes the tissues transparent and difficult to see and handle.

### 3.2 Combined HCR and IHC

Users may wish to detect mRNA expression while probing protein expression by immunohistochemistry (IHC). To demonstrate the use of *in situ* HCR with IHC, we probed *Antp*, *Ubx*, and *Abd-B* mRNA expression individually with antibodies directed to the proteins of each gene. When combining *in situ* HCR with IHC, the HCR protocol is performed as described above, but instead of mounting, users proceed with the IHC protocol described below. The combined HCR/IHC protocol takes five days to complete. An overview of the protocol workflow is illustrated in **Fig. 1**. The results of these experiments are shown with exemplary images presented in **Fig. 3**. Although we apply the IHC protocol after the *in situ* HCR module, users may try performing IHC before *in situ* HCR, in case immunostaining quality is compromised from the HCR procedure.

**Figure 3.**
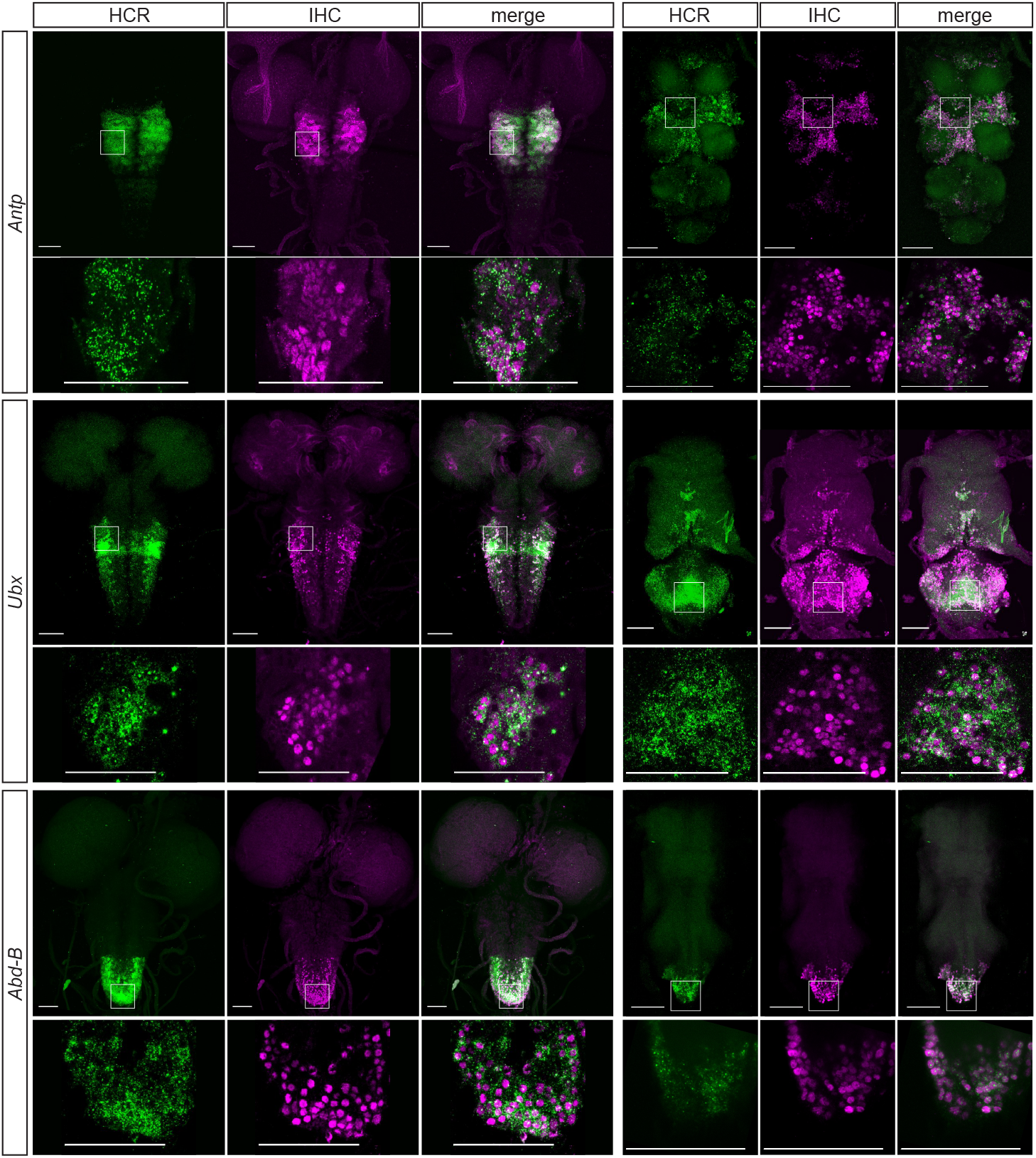
Combined *in situ* HCR and immunostaining to detect *Hox* mRNA and protein expression in the *D. melanogaster* larval and adult central nervous system. Antp-B1, Ubx-B2, and Abd-B-B3 probe-sets were used individually with Alexa Fluor® 488-based amplifiers to detect *Hox* mRNA expression. The *in situ* HCR protocol was followed with IHC using antibodies against individual Hox proteins. As expected, mRNA and protein expression of each *Hox* gene occurs in largely the same cells in the larval and adult central nervous system. While Hox protein expression is nuclear, mRNA expression is largely cytoplasmic. Scale bar = 50 μm.

#### Day 3

1. A post-HCR PFA fixation may be applied to stabilize HCR signals. Note that post-HCR fixation may affect IHC signals depending on the antibody. Fixation conditions, including PFA concentration and fixation time, can be adjusted to determine the conditions to obtain optimal HCR and IHC results. Here, we perform 2% PFA fixation for 35 min at RT for adult CNS and no fixation for larval CNS. The nervous systems are rinsed three times and washed for 20 minutes in PBTx before applying the fixative or moving onto the following steps.
2. Remove the fixative (if applied), and rinse samples three 3 times in PBTx at RT. Incubate the samples in a blocking solution (*e.g*., 5% NGS in PBTx) for 1.5 hours at room temperature.
3. Replace the blocking solution with a primary antibody prepared in blocking solution, cover and seal the dish with parafilm, and incubate overnight at 4°C.

#### Day 4

4. Remove the primary antibody solution and rinse the nervous systems three times in PBTx. Then wash three times, 30 minutes each, with PBTx.
5. Replace the final wash with the secondary antibody prepared in blocking solution, cover and seal the dish with parafilm, and incubate overnight at 4°C.

#### Day 5

6. Remove the secondary antibody solution and rinse the nervous systems three times in PBTx. Then wash three times, 30 minutes each, with PBTx.
7. Proceed with the mounting protocol.

### 3.3 Applications: Comparing gene expression in central nervous system of Drosophila species

Like other animals, species in the genus *Drosophila* exhibit a rich diversity of behaviors, which likely evolved through changes in the expression of genes that influence the morphology, physiology, or molecular properties of discrete neurons in the nervous system. The protocol we describe in this chapter provides an effective way to visualize gene expression in the central nervous system across the *Drosophila* phylogeny and identify gene expression changes that are relevant to the evolution of nervous systems and behaviors.

To demonstrate this application, we visualized the mRNA expression of *Antp*, *Ubx*, and *Abd-B* in the *D. yakuba* nervous system, a closely related species that diverged from *D. melanogaster* ~12.8 million years ago [10]. Using the protocol described above with the initiators and amplifier hairpins we applied to *D. melanogaster* nervous systems, we probed the expression of *Antp*, *Ubx*, and *Abd-B* in the *D. yakuba* larval and adult ventral nerve cords. The expression of all three genes in the *D. yakuba* larval and adult nervous system (**Fig. 4**) was detected in a pattern that was similar to that observed in the *D. melanogaster* nervous system (**Fig. 2**). The sensitivity of detection using *D. yakuba* nervous systems was comparable to that obtained using *D. melanogaster* nervous systems. This suggests that when qualitatively comparing mRNA expression of well-conserved orthologous genes in the nervous systems across *Drosophila* species, users can potentially apply a single common probe-set.

**Figure 4.**
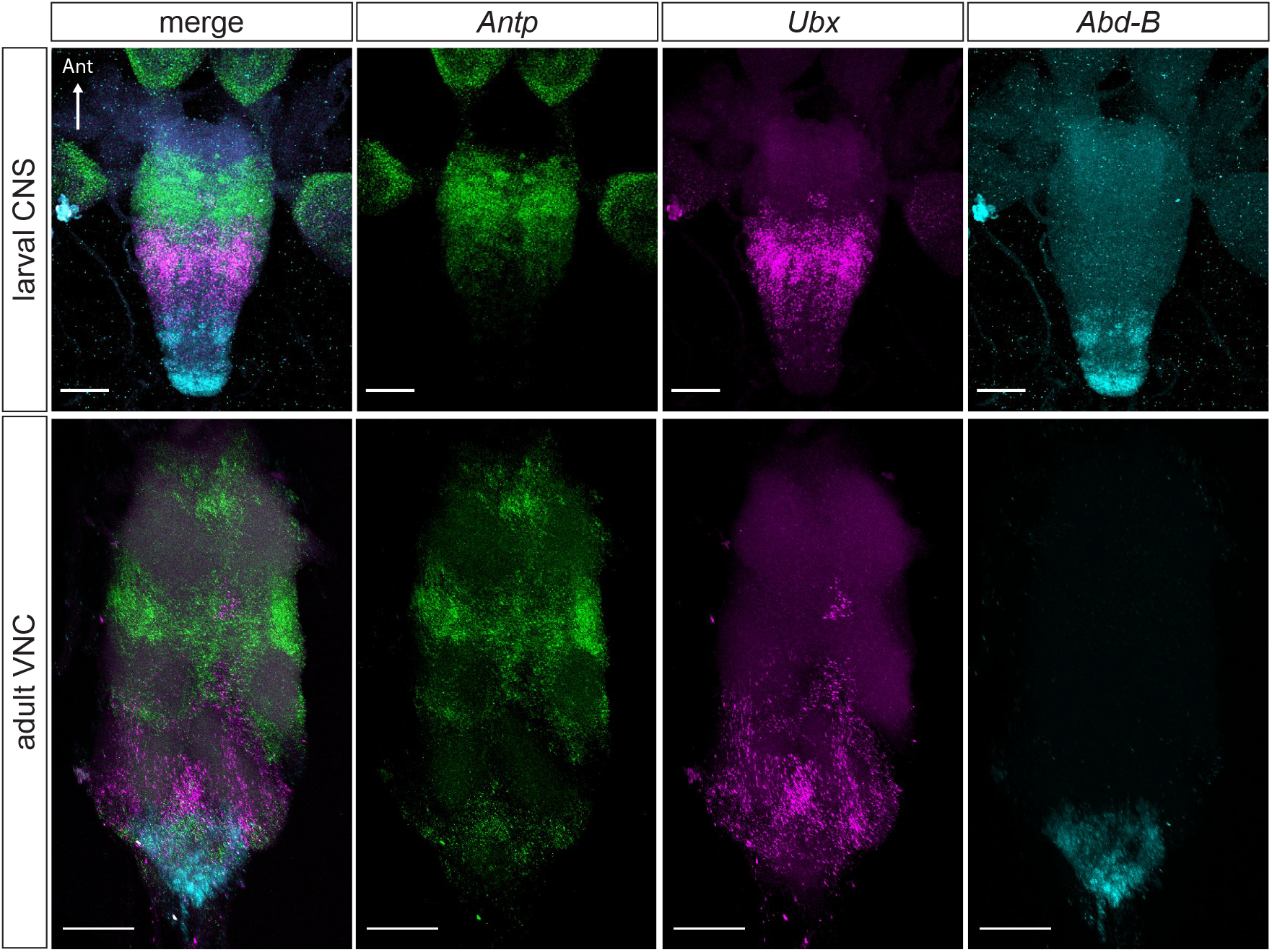
Detection of *Hox* gene mRNA expression in the larval and adult central nervous system of *D. yakuba* by *in situ* HCR. Using the same probe-sets and amplifiers as those used in the experiment shown in Fig. 1, *Hox* mRNA expression was detected in the larval and adult central nervous system of *D. yakuba*. *Antp*, *Ubx*, and *Abd-B* expression in the *D. yakuba* nervous system is similar to that observed in *D. melanogaster*. Scale bar = 50 μm.

### 3.4 Applications: Validation of single-cell RNA-Seq results

Recent advances in single-cell mRNA sequencing technologies (scRNA-Seq) have been transformative in generating atlases of gene expression with cellular resolution and identifying neuronal cell types defined by expression of unique molecular markers [11]. Fluorescence *in situ* hybridization provides an effective means to visualize the expression of genes identified in scRNA-Seq experiments, thus providing a powerful way to validate identified molecular markers; identify homologous populations of neurons across conditions and genotypes; and guide subsequent studies to determine if and how a given candidate gene influences neural development, physiology, and behavior.

Most neurons in the adult *Drosophila* central nervous system are born during larval life from a set of segmentally repeating, bilateral neural stem cells called neuroblasts [12]. A given neuroblast will often give rise to two lineages, *i.e*., hemilineages, of neurons. Neurons of a given hemilineage may share molecular, anatomical, or functional properties [12–15]. A recent study used a scRNA-Seq approach to identify new molecular markers that label neurons derived from one or more hemilineages in the *Drosophila* ventral nerve cord [9]. One marker, *Sox21B*, was found to label ventral nerve cord neurons specifically from hemilineage 13B, but no antibody against SOX21B protein is available for validation. To demonstrate the application of our protocol in validating scRNA-Seq data, we used *in situ* HCR to visualize the expression of *Sox21B* mRNA in the adult and larval central nervous system. Consistent with published RNA-Seq data, *Sox21B* expression was detected in a cluster of segmentally repeating cells located ventromedially in the three thoracic neuromeres of the adult ventral nerve cord (**Fig. 5**). These clusters are situated where neurons derived from hemilineage 13B are normally found [12]. *Sox21B* expression was also detected in segmentally repeating ventromedial groups of cells in the larval nerve cord, which may correspond to neurons of hemilineage 13B (**Fig. 5**). In addition, *Sox21B* was widely expressed in the adult brain, consistent with published scRNA-Seq data of the *Drosophila* brain [16]. In the larval and adult nervous systems, *Sox21B* was found in several bilateral clusters of cells outside of hemilineage 13B (**Fig. 5**).

**Figure 5.**
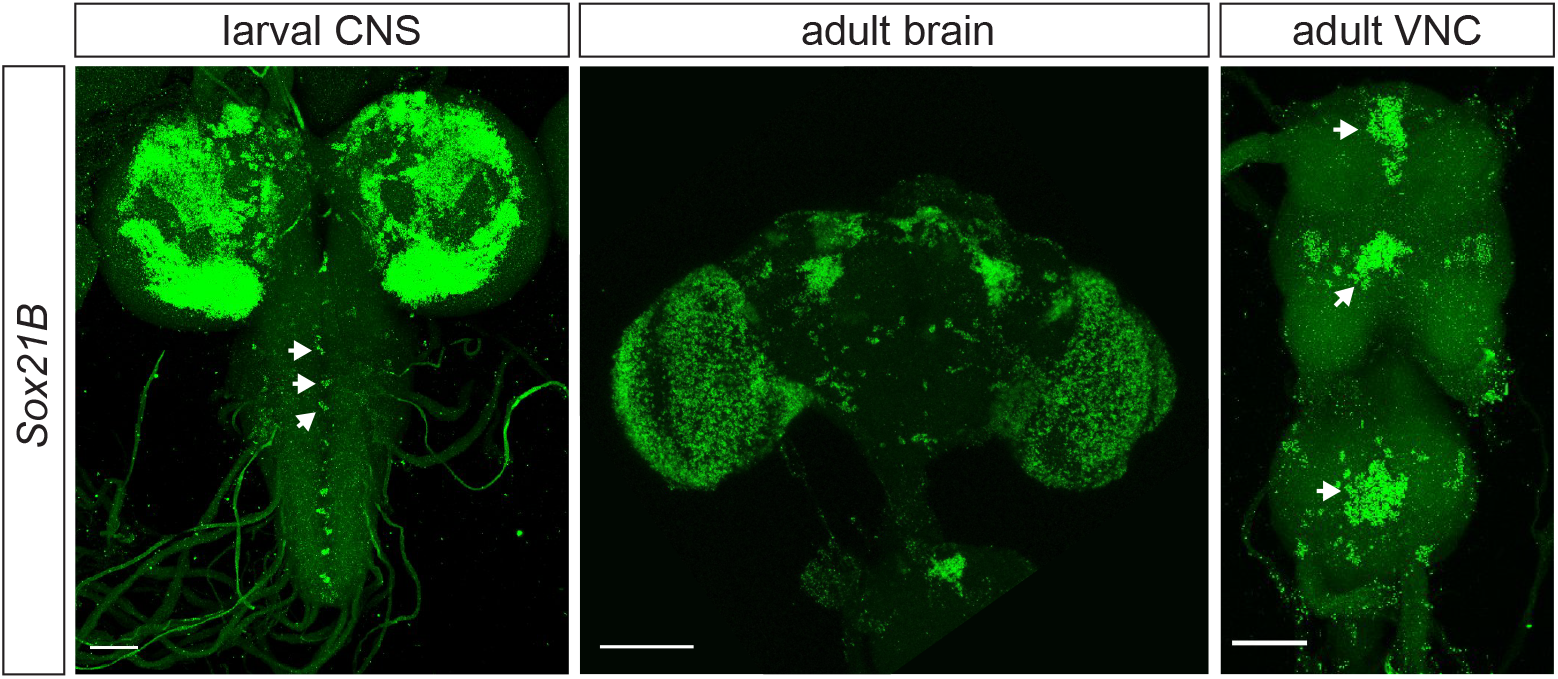
Validation of scRNA-Seq results using *in situ* HCR. *Sox21B* expression in the larval and adult central nervous system of *D. melanogaster* was detected by *in situ* HCR. In a previous study (Allen *et al*., 2020), expression of *Sox21B* was found in neurons derived from hemilineage 13B using scRNA-Seq. Expression of *Sox21B* in hemilineage 13B was visualized by *in situ* HCR using a B1 probe-set against *Sox21B* and a B1-Alexa Fluor® 488 amplifier. Neurons of hemilineage 13B occur as segmentally repeating ventromedial clusters through the thoracic neuromeres of the ventral nerve cord. *Sox21B* expression occurs in clusters of cells where hemilineage 13B normally resides in the adult and larval ventral nerve cord (arrowheads). *Sox21B* is also expressed in cells of the larval and adult brain. Scale bar = 50 μm.

## 4. NOTES

The protocols presented in this chapter provide an effective, relatively affordable and straightforward means to visualize mRNA and protein expression in whole-mount adult and larval central nervous systems. The protocol is especially useful for applications where antibodies are often not available or ideal such as cross-species gene expression comparisons and validation of RNA-Seq results in the fly nervous system. Although our protocol focuses on quantitative mRNA detection with subcellular resolution, *in situ* HCR is also compatible with single-molecule imaging. For such applications, Molecular Instruments™ recommends the use of relatively large probe sets (*e.g*., probe-sets of ≥30) for maximal signal-to-noise fidelity, and shorter amplification times (*e.g*., 45–90 min) for short amplification polymers that allow individual mRNAs to be resolved as diffraction-limited dots [2].

Based on our experiences, we share a few notes below for users to further optimize the protocol according to their specific needs.

### 4.1 Sample Treatment: Proteinase K digestion and post-hybridization paraformaldehyde fixation

Proteinase K digestion is commonly used in *in situ* protocols as a way to increase the permeability of thick whole-mount tissues and enhance *in situ* probe penetration. Using the experimental conditions described above, the addition of a Proteinase K step did not offer any obvious benefit for adult or larval nervous systems. Applying Proteinase K at a concentration of 4 μg/ml for 7 minutes after fixation and before hybridization had no noticeable effect on the signal-to-noise ratio of the mRNAs we detected (data not shown). Nevertheless, users should keep in mind that optimized Proteinase K treatment can, in some circumstances, improve *in situ* results, whether due to better tissue penetration or to removal of RNA-binding proteins that thwart probe hybridization (H. Choi, personal communication).

Prior to immunostaining, a post-hybridization paraformaldehyde fixation step may be applied to stabilize the amplification polymers and HCR signals. We found that this treatment appeared to compromise immunostaining quality in an antibody-specific manner. For instance, some antibodies, such as rabbit anti-GFP (ThermoFisher, A-11122), were largely insensitive to post-HCR fixation; however other antibodies, such as anti-N-Cadherin (DSHB, BP104), were highly sensitive to it (data not shown). We have tested three post-HCR fixation conditions (*i.e*., no fix, 2% paraformaldehyde, or 4% paraformaldehyde each for 35 minutes) and found that the absence of a post-HCR fixation step performs relatively better for larval nervous tissues, while a 2% post-HCR fixation performs better for the adult (data not shown). We recommend that users test different post-HCR fixation conditions to achieve their desirable balance between *in situ* and immunostaining signals. Furthermore, users can try IHC before *in situ* HCR, if post-HCR immunostaining is consistently problematic.

### 4.2 Detection sensitivity: probe concentration, probe-set size, and amplification conditions

*In situ* HCR is in principle well suited to detect low abundance transcripts. In other mRNA-FISH approaches for whole-mount *Drosophila* nervous systems [5, 6], the fluorophores are directly conjugated to each individual probe; *e.g*., a probe-set of 20 will lead to the assembly of, at most, 20 fluorophores per transcript. It is estimated that most amplification polymers in *in situ* HCR will each contain ~200 fluorophores [3]. Remarkably, this means that a set of 20 probes could, in theory, lead to the construction of upwards of 4000 fluorophores on each transcript.

If users are probing the expression of a low abundance target gene, there are some “tweaks” that may improve detection sensitivity. First, users can try adjusting the concentration of the probe. The standard probe concentration, as recommended by Molecular Instruments™, is 4 nM, but increasing probe concentration may improve detection sensitivity. This adjustment can be effective and cost-free when users have an excess of probes. Second, the size of the probe-set can be increased. In our experiments, we used a set of 20 split-initiator probe pairs, each targeting a total of 52 bases within an mRNA type (*i.e*., 26 bases for each split-initiator probe of a pair). Using our protocol, a probe-set size of 20 consistently yielded reliable signals for a variety of target genes. However, increasing the probe-set size to 30 or more may further enhance detection, especially for transcripts with low levels of gene expression. Third, confocaling under higher magnification can improve the signal-to-noise ratio. Finally, adjusting the amplification conditions may also enhance sensitivity. In our experiments, we typically used an amplification time of 16 hours. Although longer amplification time beyond overnight is not expected to strengthen HCR signals, increasing the hairpin concentration may help (H. Choi, personal communication).

### 4.3 Cost

Another major advantage of *in situ* HCR over other mRNA-FISH protocols [5, 6] is in the cost. Currently, other published mRNA-FISH protocols on whole-mount *Drosophila* nervous systems [5, 6] utilize probe-sets supplied by Stellaris™, which start at US$675 and provide enough material for 200–400 experiments [6]. At the time of writing this chapter, a set of 20 customized HCR probe pairs starts at US$252 for academic and non-profit users, which allows ~250 *in situs*. With the amplifier mix currently starting at US$97.50, the cost of probing the mRNA expression of a single gene is notably more affordable using HCR than using Stellaris-based approaches. The savings in cost are more substantial if users wish to detect the expression of multiple genes, as a single amplifier mix can be used with multiple HCR probe-sets. Furthermore, if one wishes to visualize transcripts using a different fluorophore, it can be easily achieved by adopting a different amplifier mix instead of purchasing a new probe-set.

## 5. ACKNOWLEDGEMENTS

We thank Dr. Lisha Shao for initially bringing the HCR method to our attention and sharing useful tips during our protocol development; Dr. Harry Choi from Molecular Instruments™ for valuable inputs; and Dr. Justin Walsh for scRNA-Seq analyses.

